# Genomic comparison of non-photosynthetic plants from the family Balanophoraceae with their photosynthetic relatives

**DOI:** 10.1101/2020.09.20.305011

**Authors:** Mikhail I. Schelkunov, Maxim S. Nuraliev, Maria D. Logacheva

## Abstract

The plant family Balanophoraceae consists entirely of species that have lost the ability to photosynthesize. Instead, they obtain nutrients by parasitizing other plants. Recent studies have shown that plastid genomes of Balanophoraceae have a number of interesting features, one of the most prominent of those being a tremendous AT content close to 90%. Also, the nucleotide substitution rate in the plastid genomes of Balanophoraceae is greater by an order of magnitude compared to photosynthetic relatives, without signs of relaxed selection. All these features have no definite explanations.

Given these unusual features, we supposed that it would be interesting to gain insight into the characteristics of *nuclear* genomes of Balanophoraceae. To do this, in the present study we analysed transcriptomes of two species from Balanophoraceae, namely *Rhopalocnemis phalloides* and *Balanophora fungosa*, and compared them with transcriptomes of their close photosynthetic relatives *Daenikera* sp., *Dendropemon caribaeus, Malania oleifera*.

The analysis showed that the AT content of nuclear genes of Balanophoraceae does not markedly differ from that of photosynthetic relatives. The nucleotide substitution rate in genes of Balanophoraceae is for an unknown reason several times larger than in genes of photosynthetic Santalales, though the negative selection in Balanophoraceae is likely stronger.

We observed an extensive loss of photosynthesis-related genes in Balanophoraceae. Also, for Balanophoraceae we did not see transcripts of several genes whose products participate in plastid genome repair. This implies their loss or very low expression, which may explain the increased nucleotide substitution rate and AT content of the plastid genomes.

## Introduction

Plants are often thought to create organic substances by photosynthesis. However, there were several dozen independent cases when a photosynthetic plant obtained an ability to get organic substances from other plants or fungi, this is called mixotrophy or partial heterotrophy (reviewed by Těšitel (2018)). Some mixotrophic plants can later lose the photosynthetic ability and start to rely solely on parasitism on their plant or fungal host, this is called complete heterotrophy, holo-heterotrophy, or just “heterotrophy” (reviewed by Merckx, Bidartondo and Hynson (2009) and by Nickrent (2020)).

This transition to heterotrophy leaves a noticeable trace on plant morphology and ecology. Heterotrophic plants usually have no leaves or very reduced ones. These plants have no to very little green color due to the absence or very low levels of chlorophyll. The majority of heterotrophic species spend most of the year completely underground, since they do not need light for survival. Instead, they appear aboveground only for reproduction. There are also other morphological and ecological changes, which have been reviewed, for example, by Leake (1994) and Těšitel (2016).

Except for some rare cases (Molina et al., 2014), plants have three genomes: the plastid, the mitochondrial and the nuclear. The plastid genome is the smallest and has the highest copy number, thus alleviating sequencing and assembly. This is the reason why most plant genomes sequenced and assembled to date are plastid ones. As the majority of plastid genes are needed for photosynthesis, it is unsurprising that one of the plastid genome changes characteristic of non-photosynthetic plants is a massive plastid gene loss (reviewed by Wicke and Naumann (2018)). Another widely observed feature is an increase in the nucleotide substitution rate, but likely not due to relaxation of natural selection. The third common feature is an increase in the AT content. While the cause of the first feature is obvious, the causes of the second and of the third still remain unknown.

The mitochondrial genomes of non-photosynthetic plants are poorly studied, but, probably, a feature that unites many of them is the horizontal transfer of genes from mitochondrial genomes of their hosts to mitochondrial genomes of parasites (Mower, Jain & Hepburn, 2012; Sanchez-Puerta et al., 2017, 2019; Roulet et al., 2020).

The nuclear genome sequencing is much more expensive than the mitochondrial or the plastid, due to the much larger size of the nuclear genome. Therefore, to obtain information about the nuclear genes, scientists often sequence the transcriptome instead of the genome itself. Various studies of nuclear genomes, including studies based on the transcriptome analysis, revealed several features characteristic of genomes of non-photosynthetic plants (Wickett et al., 2011; Lee et al., 2016; Schelkunov, Penin & Logacheva, 2018; Yuan et al., 2018). First, just as in the plastid genomes, there is an extensive loss of photosynthesis-related genes. However, for a yet-unknown reason, genes participating in chlorophyll synthesis are often retained. It is hypothesized that the chlorophyll may have some other function apart from photosynthesis. Just as in plastid genomes, for an unknown reason the nuclear genes of non-photosynthetic plants evolve faster and without signs of relaxed selection. The AT content of nuclear genes of non-photosynthetic plants is not much increased or not increased at all, unlike to the plastid genomes.

Among the most unusual plastid genomes currently known to scientists are plastid genomes of non-photosynthetic plants from the family Balanophoraceae (Su et al., 2019; Schelkunov, Nuraliev & Logacheva, 2019; Chen et al., 2020a). The family Balanophoraceae comprises several dozen species that inhabit tropical and subtropical areas and feed by attaching to roots of different plants and sucking nutrients from them. Plastid genomes have been sequenced for one species of the genus *Rhopalocnemis* and four species of the genus *Balanophora*. These plastid genomes are about 10 times smaller than plastid genomes of typical photosynthetic plants. The AT content in currently known plastid genomes of Balanophoraceae is in the range of 86.8%-88.4%, which makes them the most AT-rich of all known plant genomes, counting not only among plastid but also among mitochondrial and nuclear genomes. This large AT content affects not only non-coding regions but also genes, with AT contents of some genes exceeding 90%. The nucleotide substitution rate in the known plastid genomes of Balanophoraceae is more than 10 times greater than in photosynthetic relatives. To top it all, at least some plastid genomes of plants from the genus *Balanophora* have changed their genetic code, with the TAG codon now coding for the tryptophan instead of being a stop-codon. As was mentioned above, such large AT contents and substitution rates currently have no accepted scientific explanation.

We supposed that nuclear genomes of plants with such unusual plastid genomes also deserve being studied. Our previous estimate for *Rhopalocnemis phalloides* suggests a nuclear genome size of at least 30 Gbp (Schelkunov, Nuraliev & Logacheva, 2019), thus nuclear genome sequencing is unpractical. Instead, we sequenced the transcriptome of *Rhopalocnemis phalloides* and also used the transcriptome of *Balanophora fungosa*, another plant from Balanophoraceae (Fig. 1), that was sequenced as part of the 1KP Project (Carpenter et al., 2019, p.). For comparison, we used the mixotrophic plants *Daenikera* sp., *Dendropemon caribaeus, Malania oleifera*. These three mixotrophic species belong to three different families of the order Santalales, where the family Balanophoraceae also belongs. The families are, according to the APG IV classification system (The Angiosperm Phylogeny Group, 2016), Olacaceae, Santalaceae and Loranthaceae respectively. Just as plants of Balanophoraceae, these three mixotrophic species are capable of parasitism, but at the same time they are obligate autotrophs, meaning that they photosynthesize and cannot survive without light (Vidal-Russell & Nickrent, 2008; Caraballo-Ortiz et al., 2017; Li, Mao & Li, 2019). Transcriptomes of these three species were sequenced earlier as parts of different projects (Xu et al., 2019; Carpenter et al., 2019).

**Figure 1.**
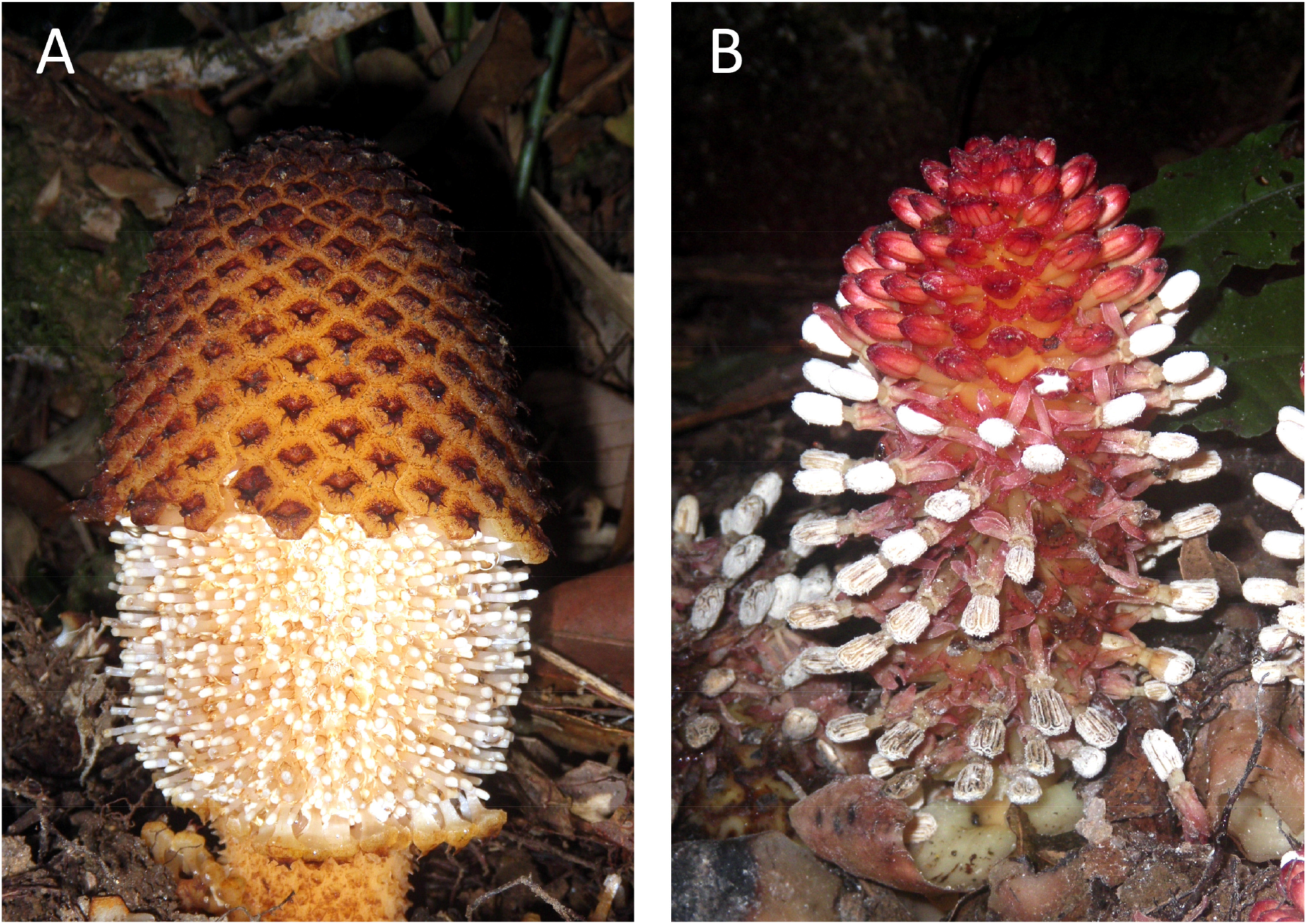
Photos of the studied non-photosynthetic species. The photos depict above-ground parts of *Rhopalocnemis phalloides* (A) and *Balanophora fungosa* (B). Photographed by M. Nuraliev.

## Materials and Methods

### Sample collection and sequencing

The sequenced specimen of *Rhopalocnemis phalloides* was collected during an expedition of the Russian-Vietnamese Tropical Centre in Kon Tum Province, Vietnam, in May 2015. The specimen was preserved in silica gel and RNAlater. The voucher is deposited at the Moscow University Herbarium (Seregin, 2018) with the barcode MW0755444.

We extracted RNA from the inflorescence. The extraction was performed using the RNEasy Mini kit (Qiagen, the Netherlands) following the manufacturer’s instructions. The only modification was the addition of Plant Isolation Aid Solution (ThermoFisher, USA) into the lysis buffer. Selection of RNA was made using the Ribo-Zero Plant Leaf kit (Illumina, USA). Further library preparation was performed with the NEBNext Ultra II RNA kit (New England Biolabs, USA).

The library was sequenced on:

1. Illumina NextSeq 500, producing 6,912,802 paired-end 75 bp-long reads (3,456,401 read pairs).
2. Illumina HiSeq 2500, producing 54,794,466 paired-end 125 bp-long reads (27,397,233 read pairs).
3. Illumina HiSeq 4000, producing 181,190,240 paired-end 76 bp-long reads (90,595,120 read pairs).

Overall, 242,897,508 reads were produced for *Rhopalocnemis phalloides* (121,448,754 read pairs).

### Transcriptomes of comparison

As transcriptomes of comparison, we used transcriptomes of *Balanophora fungosa, Daenikera* sp., *Dendropemon caribaeus* and *Malania oleifera. Balanophora fungosa, Daenikera* sp. and *Dendropemon caribaeus* were sequenced as part of the 1KP project (Carpenter et al., 2019), while *Malania oleifera* was sequenced by Xu et al. (2019).

The transcriptomes of *Balanophora fungosa, Daenikera* sp., *Dendropemon caribaeus* and *Malania oleifera* were sequenced on Illumina HiSeq 2000, producing:

1. For *Balanophora fungosa* 26,470,118 paired-end 90 bp-long reads (13,235,059 read pairs). The identifier for these reads in the NCBI Sequence Read Archive (SRA) is ERR2040275.
2. For *Daenikera* sp. 22,878,728 paired-end 90 bp-long reads (11,439,364 read pairs). The SRA identifier of reads is ERR3487343.
3. For *Dendropemon caribaeus* 22,393,054 paired-end 90 bp-long reads (11,196,527 read pairs). The SRA identifier of reads is ERR2040277.
4. For *Malania oleifera* 222,952,532 paired-end 150 bp-long reads (111,476,266 read pairs). The SRA identifiers of reads are SRR7221530, SRR7221531, SRR7221535, SRR7221536, SRR7221537.

To minimize methodological differences, we assembled the transcriptomes of these 4 species with exactly the same methods as the transcriptome of *Rhopalocnemis phalloides* (see below). For *Rhopalocnemis phalloides* and *Malania oleifera* which have several read datasets, the datasets were combined before assembly.

Initially we used *Ximenia americana* (SRA identifier ERR2040276) instead of *Malania oleifera*, but the BUSCO analysis indicated a poor assembly quality, thus it was replaced by *Malania oleifera*.

### Read processing and transcriptome assembly

Reads were trimmed by Trimmomatic 0.39 (Bolger, Lohse & Usadel, 2014), performing 5 procedures in the following order:

1. Adapters was trimmed by the palindromic method.
2. Bases with sequencing quality less than 3 were trimmed from the 3’ end.
3. If a group of 4 consecutive bases with an average quality of less than 15 existed, this group was trimmed together with the part of the read that was in the 3’ direction from that group.
4. If the average quality of the read was less than 20, the read was discarded entirely.
5. If the length of the read after the previous 4 steps was less than 30 bases, it was removed.

The assembly was made by Trinity 2.8.4 (Haas et al., 2013) with digital normalization to 50× coverage. The minimum contig length was set to 200 bp. The expression levels of the assembled transcripts were quantified by Salmon 0.9.0 (Patro et al., 2017). As the major isoform for a gene we defined the isoform with the highest expression level, measured by the “transcripts per million” (TPM) value. The minor isoforms were discarded.

Contigs with very low coverage are likely to contain misassemblies. To detect such contigs, the reads were mapped to all contigs by CUSHAW 3.0.3 (Liu, Popp & Schmidt, 2014) requiring at least 80% of a read to map with a sequence similarity of at least 98%. Contigs with an average coverage less than 3× were discarded.

Protein-coding sequences (CDSs) were predicted by TransDecoder 5.5.0 (Haas et al., 2013), using its in-build capabilities and also using the presence of Pfam-A domains in open reading frames (ORFs) and the similarity of ORF sequences to sequences in the NCBI NR database. The NCBI NR database was current as of 13th May 2019. The Pfam-A domain prediction in ORFs found by TransDecoder was conducted using the Pfam-A 32.0 database (El-Gebali et al., 2019) and the hmmscan tool from the HMMER 3.2 package (Mistry et al., 2013) with the default parameters. The similarity search between ORFs and NCBI NR sequences was performed by the “blastp” command from DIAMOND 0.9.25 (Buchfink, Xie & Huson, 2015) with the “--more-sensitive” option and a maximum e-value of 10^−5^. An ORF was considered a probable CDS if at least one of the three criteria was met:

1. TransDecoder considered this ORF a likely CDS based on its in-built criteria, like the hexanucleotide frequencies.
2. A Pfam-A domain was found in the ORF.
3. The ORF has a match in NCBI NR.

After the CDS prediction, we removed CDSs with proteins having best matches in NCBI NR not to Embryophyta, thus removing contamination. By proteins we hereafter mean translated CDSs. CDSs with proteins that had no matches in NCBI NR, and consequently no definite taxonomic assignment, were kept.

The completeness of the transcriptome assemblies was assessed by BUSCO 3.1.0 (Simão et al., 2015) using the eukaryotic set of conserved proteins. The eukaryotic set was preferred over plant sets because plant sets contain some proteins that function in photosynthesis and thus should be absent in *Rhopalocnemis phalloides* and *Balanophora fungosa*.

### Transcriptome annotation

The transcriptome was annotated using the following three methods:

1. The method of reciprocal best hits (RBH). Major isoforms of proteins of *Arabidopsis thaliana* from the TAIR10 database (Berardini et al., 2015) were aligned by BLASTP 2.9.0+ (Camacho et al., 2009) to proteins of each of the 5 studied species with the maximum e-value set to 10^−5^, word size 3 and the low-complexity filter switched off. Then, vice versa, the proteins of the 5 species were aligned in the same way to the proteins of *Arabidopsis thaliana*. If a pair of proteins were reciprocal best hits (or, which is the same, “reciprocal best matches”) to each other in both of these alignments, the corresponding protein in the studied species was supposed to have the same functions as its match in *Arabidopsis*.
2. The Gene Ontology (GO) annotation (The Gene Ontology Consortium, 2019). Proteins of the 5 studied species were annotated with GO terms by PANNZER2 (Törönen, Medlar & Holm, 2018) using the positive predictive value threshold of 0.5. The relaxed criteria for query and subject lengths were switched on by the option PANZ_FILTER_PERMISSIVE”.
3. The KEGG metabolic annotation (Kanehisa et al., 2016). The annotation was performed by the GhostKOALA (Kanehisa, Sato & Morishima, 2016) web server on 24th December 2019.

The second and the third methods were used only for nuclear proteins of the studied species. To remove transcripts of mitochondrial and plastid proteins, we removed all transcripts whose proteins were RBHs to mitochondrial and plastid proteins of *Arabidopsis thaliana* in the first analysis.

### Phylogenetic tree and the natural selection analysis

To construct the phylogenetic tree of the 5 studied species, we determined orthogroups of their CDSs and CDSs of *Arabidopsis thaliana*. The CDSs of *Arabidopsis thaliana* used for this analysis were CDSs from the major transcript isoforms from the TAIR10 database. To restrict the analysis to the nuclear CDSs, the mitochondrial and plastid CDSs were excluded as described above. The orthology was determined by OrthoFinder 2.3.8 (Emms & Kelly, 2019), using DIAMOND as a tool for similarity calculation.

Then, to build the tree, we took all orthogroups that had exactly one sequence from each species, there were 1039 such orthogroups. They were aligned by TranslatorX 1.1 (Abascal, Zardoya & Telford, 2010) with MAFFT 7.402 (Katoh & Standley, 2013). The source code of TranslatorX was changed to use MAFFT in the E-INS-i mode which allows for large gaps in the alignment. This is useful, because the exonic content can be different in different orthologs. The poorly aligned regions of orthogroups were removed by Gblocks 0.91b (Castresana, 2000) with the default parameters. Then the alignments for different orthogroups were concatenated into one alignment. The tree for this alignment was built by RAxML 8.2.12 (Stamatakis, 2014) with the GTR+Gamma model, using 20 starting trees and with the number of bootstrap pseudoreplicates automatically determined by the autoMRE method.

The selection analysis was performed by PAML 4.9 (Yang, 2007) using the branch model with the F3×4 codon frequencies model, the starting dN/dS of 0.5 and the starting transition/transversion ratio of 2. The option “cleandata”, which removes columns with gaps and stop codons, was switched on. The tree given to PAML was the tree produced by RAxML. The numbers of substitutions that happened on tree branches were calculated by PAML. To calculate 95% confidence intervals for dN/dS values of branches we generated 1000 bootstrap pseudoreplicates for the alignment, performed 1000 separate PAML calculations and then took 2.5th and 97.5th percentiles for dN/dS on each branch.

The tree was drawn by TreeGraph 2.14.0 (Stöver & Müller, 2010) with *Arabidopsis thaliana* used as the outgroup. As the number of substitutions on branches we used values provided by PAML, not RAxML.

AT contents of nuclear genes were calculated from the same concatenated gene alignment as was used for the phylogenetic tree construction and the dN/dS evaluation. dN/dS values for the *NTH1* gene were computed by the same method as the dN/dS values of concatenated genes, using the topology inferred from the concatenated genes.

### Other analyses

The GO enrichment analysis was performed only for nuclear genes, excluding plastid and mitochondrial genes as described above. For each GO term we calculated the p-value for the difference in the proportion of genes that code for this GO term between each of the 2 nonphotosynthetic species and 3 photosynthetic species, thus making 6 comparisons overall. The p-values were calculated through the Fisher’s exact test by the program GOAtools 0.6.10 (Klopfenstein et al., 2018). Then we performed the Bonferroni correction for these 6 comparisons and then the Benjamini-Hochberg correction for all GO terms. Differences between photosynthetic and non-photosynthetic species that had q-values less than or equal to 0.05 were considered statistically significant.

Information about the plastid gene content in *Rhopalocnemis phalloides* and *Balanophora fungosa* was obtained from the papers that described their plastid genomes (Schelkunov, Nuraliev & Logacheva, 2019; Chen et al., 2020a). The plastid genome sequences of *Daenikera* sp. and *Dendropemon caribaeus* are still unknown. However, given the similar gene content in the plastid genome of photosynthetic Santalales (Chen et al., 2020a), it is reasonable to assume that the plastid gene content in *Daenikera* sp. and *Dendropemon caribaeus* is approximately the same. There are two scientific papers about the plastid genome of *Malania oleifera*, which highly contradict each other (Yang & He, 2019; Xu et al., 2019). One of them states that the genome length is 158,163 bp, while the other tells about a 125,050 bp-long genome. Given that the gene content in the second genome highly differs from gene contents in other photosynthetic Santalales (Chen et al., 2020a), we suppose that the second genome either contains assembly mistakes or belongs to some other species, misidentified as *Malania oleifera*. Therefore, as the true plastid genome of *Malania oleifera* we consider the longer one.

## Results and Discussion

### The assemblies

Results of the BUSCO analysis for the assemblies are provided in Fig. 2. The results for all transcripts indicate the near absence of missing genes. However, at the stage of minor isoforms removal, the percentage of missing and fragmented BUSCO genes increased. This is probably because as “the major isoforms” we defined the most expressed isoforms, and these most expressed isoforms may not have the same exons that BUSCO gene models have. Such a difference in exonic content may make BUSCO underestimate the completeness. More mechanistic measures, such as N50 and the total numbers of contigs in the assemblies are provided in Table S1.

**Figure 2.**
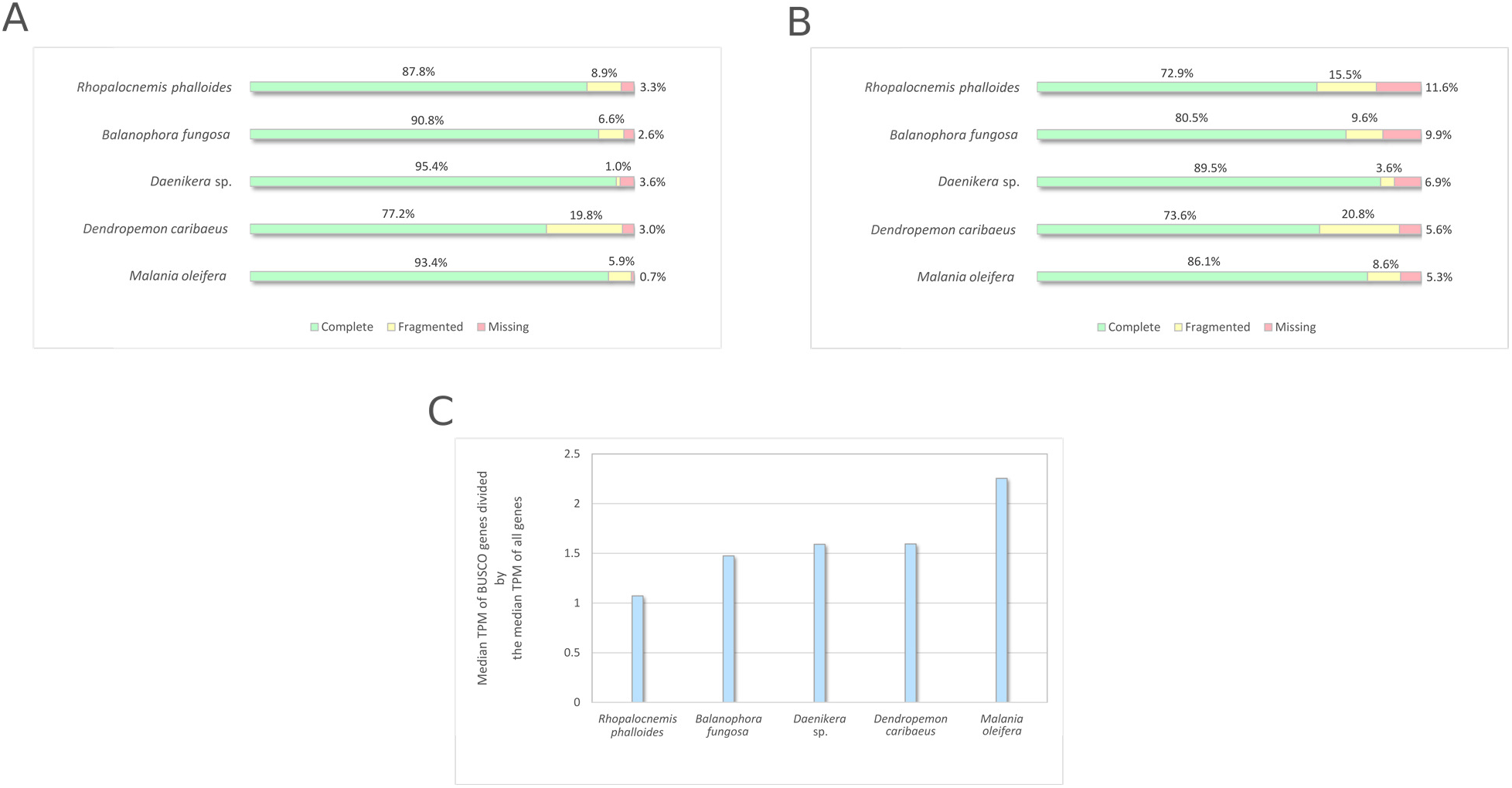
BUSCO analysis results. (A) BUSCO results for all sequences produced by Trinity. (B) BUSCO results for CDSs from major isoforms of transcripts after removal of transcripts with low coverage and contamination. (C) Comparison of expression of BUSCO genes and all genes.

It should be noted that the BUSCO analysis may be less suitable to assess the completeness of transcriptome assemblies compared with genome assemblies if BUSCO genes are expressed at higher levels than average genes. A comparison of the median expression level of BUSCO genes with the median expression level of all genes (Fig. 2c) suggests that the median expression level of BUSCO genes is approximately 1.5 times larger. In this comparison as “all genes” we used the same CDS set as in Fig. 2b, but omitted CDSs that have no Gene Ontology (GO) terms to reduce the number of false-positively predicted genes. This difference of expression between BUSCO genes and all genes suggests that BUSCO genes may be assembled slightly better, thus the completeness of the assembly may be overestimated.

### Nuclear gene content as inferred from the transcriptome assemblies

Results of the Gene Ontology terms enrichment analysis (hereafter, “GO enrichment”) indicate that all the most statistically significant reductions in the gene sets of non-photosynthetic plants are unambiguously linked to the loss of photosynthesis.

The lists of 10 GO terms most underrepresented and most overrepresented in nonphotosynthetic plants compared to photosynthetic plants are provided in Tables 1 and 2. The complete table with all GO terms with significantly different amounts of genes is Table S2.

**Table 1.** GO terms most underrepresented among nuclear genes of non-photosynthetic Santalales compared to photosynthetic Santalales. Values in the cells are the number of genes with this GO term divided by the total number of genes with GO terms in this species.

| GO term | *Rhopalocnemis phalloides* | *Balanophora fungosa* | *Daenikera* sp. | *Dendropemon caribaeus* | *Malania oleifera* |
| --- | --- | --- | --- | --- | --- |
| photosystem I | 1/22583 | 0/12178 | 28/16391 | 23/22802 | 50/21755 |
| photosystem II | 1/22583 | 2/12178 | 41/16391 | 36/22802 | 62/21755 |
| photosystem | 3/22583 | 3/12178 | 57/16391 | 54/22802 | 86/21755 |
| chlorophyll metabolic process | 2/22583 | 2/12178 | 31/16391 | 32/22802 | 27/21755 |
| photosynthesis | 9/22583 | 6/12178 | 94/16391 | 98/22802 | 133/21755 |
| photosynthesis, light reaction | 7/22583 | 4/12178 | 32/16391 | 51/22802 | 61/21755 |
| thylakoid part | 19/22583 | 17/12178 | 118/16391 | 145/22802 | 172/21755 |
| photosynthetic membrane | 17/22583 | 16/12178 | 106/16391 | 129/22802 | 160/21755 |
| thylakoid | 17/22583 | 22/12178 | 125/16391 | 154/22802 | 189/21755 |
| thylakoid membrane | 14/22583 | 14/12178 | 87/16391 | 109/22802 | 130/21755 |

**Table 2.** GO terms most overrepresented among nuclear genes in non-photosynthetic Santalales compared to photosynthetic Santalales. Values in the cells are the number of genes with this GO term divided by the total number of genes with GO terms in this species.

| GO term | *Rhopalocnemis phalloides* | *Balanophora fungosa* | *Daenikera* sp. | *Dendropemon caribaeus* | *Malania oleifera* |
| --- | --- | --- | --- | --- | --- |
| DNA integration | 9486/22583 | 2001/12178 | 475/16391 | 344/22802 | 1714/21755 |
| DNA metabolic process | 10265/22583 | 2470/12178 | 968/16391 | 1095/22802 | 2648/21755 |
| nucleic acid metabolic process | 11374/22583 | 3449/12178 | 2266/16391 | 2967/22802 | 4308/21755 |
| nucleobase-containing compound metabolic process | 11604/22583 | 3617/12178 | 2534/16391 | 3368/22802 | 4643/21755 |
| heterocycle metabolic process | 11720/22583 | 3731/12178 | 2717/16391 | 3583/22802 | 4830/21755 |
| cellular aromatic compound metabolic process | 11735/22583 | 3733/12178 | 2745/16391 | 3619/22802 | 4881/21755 |
| organic cyclic compound metabolic process | 11783/22583 | 3777/12178 | 2799/16391 | 3694/22802 | 4942/21755 |
| nucleic acid binding | 8244/22583 | 2887/12178 | 2000/16391 | 2819/22802 | 3591/21755 |
| cellular nitrogen compound metabolic process | 11904/22583 | 3868/12178 | 2939/16391 | 3872/22802 | 5105/21755 |
| cellular macromolecule metabolic process | 12483/22583 | 4221/12178 | 3922/16391 | 5510/22802 | 6150/21755 |

As can be seen from Table 1, photosynthesis-related genes are nearly absent from the nonphotosynthetic species. The GO annotation process is known to produce a number of falsepositive results (Zhou et al., 2019), therefore the real reduction in the number of photosynthesis-related genes is likely stronger than seems from the table.

Table 2 demonstrates that the non-photosynthetic species have increased proportions of genes with the GO term “DNA integration” and some nucleotide metabolism-related GO terms. A BLAST analysis of genes with these GO terms indicates that they mostly belong to transposons. Enrichment by these GO terms was previously observed in non-photosynthetic plants of the genera *Epipogium* and *Hypopitys* (Schelkunov, Penin & Logacheva, 2018). We do not have an obvious explanation for this phenomenon.

As an alternative method to check for changes in the gene content, we composed a list of *Arabidopsis thaliana* proteins with functions interesting to us and searched for reciprocal best hits (RBHs) in the CDS sets of the five studied transcriptomes (Table S3). This analysis indicates a vast disappearance of genes linked to photosynthesis. Some genes with products participating in the Calvin cycle are retained, but these genes also have functions beyond the Calvin cycle (Schelkunov, Penin & Logacheva, 2018). A feature which deserves mentioning is the presence in non-photosynthetic species of several genes with products required for import into thylakoids. Probably *Rhopalocnemis phalloides* and *Balanophora fungosa* still have thylakoids that have functions unrelated to photosynthesis, or the products of these genes have functions beyond import into thylakoids.

An unexpected feature of many non-photosynthetic plants is presence of some levels of chlorophyll (Cummings & Welschmeyer, 1998). It was hypothesised that the chlorophyll may have some functions other than photosynthesis (Cummings & Welschmeyer, 1998). Analyses of transcriptomes and genomes showed that the pathways for synthesis and breakdown of the chlorophyll are indeed probably retained in some non-photosynthetic species (Wickett et al., 2011; Schelkunov, Penin & Logacheva, 2018; Marcin et al., 2020), though likely lost in some others (Ng et al., 2018; Schelkunov, Penin & Logacheva, 2018; Chen et al., 2020b). In *Rhopalocnemis phalloides* and *Balanophora fungosa* the RBH analysis indicates complete disappearance of chlorophyll synthesis and breakdown genes.

The plastid genomes of *Rhopalocnemis phalloides* and *Balanophora fungosa* were previously shown to be approximately 10 times shorter than plastid genomes of their close photosynthetic relatives. The number of genes in these two non-photosynthetic species is also reduced approximately tenfold. The RBH analysis revealed no transcripts of the lost genes, implying that they were not transferred to the nuclear or the mitochondrial genome, but instead disappeared completely.

As the third method of analysis of gene loss, besides GO enrichment and RBH analysis, we used KEGG metabolic annotation. Visual inspection of pathway maps produced by KEGG suggests, in addition to the conclusions that followed from the GO enrichment and RBH, that *Rhopalocnemis phalloides* and *Balanophora fungosa* lack transcripts for a number of genes with products participating in: circadian rhythm organization (Fig. S1), terpenoid synthesis (Figs. S2 and S3) and carotenoid synthesis (Fig. S4). These changes are likely linked to the loss of photosynthesis.

### AT content in nuclear genes

It was previously shown that plastid genomes of Balanophoraceae are very AT-rich, with AT contents of currently sequenced genomes being in the range of 86.8%-88.4% (Su et al., 2019; Schelkunov, Nuraliev & Logacheva, 2019; Chen et al., 2020a). This makes them the most AT-rich of any plant genomes, and one of the most AT-rich among all genomes. The AT contents of their relatives from photosynthetic families of the same order Santalales are much lower, being in the range of 61.8%-65.13%. The increased AT content of Balanophoraceae is a feature not only of non-coding regions, but of coding too, with average weighted by gene length AT contents of plastid protein-coding genes in species of Balanophoraceae being 88.1%-91.3%, and in photosynthetic Santalales being 60.9%-63.7%. The cause of such high AT contents in plastid genomes of Balanophoraceae is unknown. Such increase is a long known but still unexplained feature of plastid genomes of non-photosynthetic plants, with Balanophoraceae being just the most extreme case (Wicke et al., 2013; Schelkunov et al., 2015; Lam, Soto Gomez & Graham, 2015; Bellot & Renner, 2015; Naumann et al., 2016; Lim et al., 2016; Logacheva et al., 2016; Roquet et al., 2016; Braukmann et al., 2017; Park, Suh & Kim, 2020).

It was logical for us to have a look at whether the same difference in AT contents exist in nuclear genes. While the plastid genome is only known for *Malania oleifera*, but not for *Daenikera* sp. and *Dendropemon caribaeus*, it safe to assume that the AT contents of plastid protein-coding genes of *Daenikera* sp. and *Dendropemon caribaeus* are somewhere in the range of 60.9%-63.7% or close to that range, because this range is based on 29 plastid genomes of photosynthetic Santalales known to date. As one may see from Table 3, the AT contents of nuclear protein-coding genes of *Rhopalocnemis phalloides* and *Balanophora fungosa* do not differ much from that of their photosynthetic relatives. The AT content of *Rhopalocnemis phalloides* is even the lowest among the species.

**Table 3.**
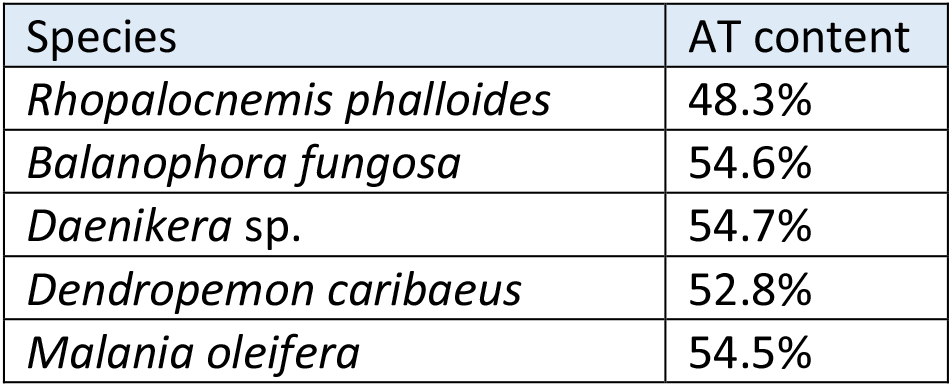
AT contents in nuclear genes of Santalales

| Species | AT content |
| --- | --- |
| *Rhopalocnemis phalloides* | 48.3% |
| *Balanophora fungosa* | 54.6% |
| *Daenikera* sp. | 54.7% |
| *Dendropemon caribaeus* | 52.8% |
| *Malania oleifera* | 54.5% |

### Nucleotide substitution rate and selection in nuclear genes

Among the unusual features shown previously for the plastid genomes of *Rhopalocnemis* and *Balanophora* are their high nucleotide substitution rates, which are approximately an order of magnitude greater than in photosynthetic relatives from the order Santalales. One may assume that this is due to relaxed natural selection. However, it was demonstrated that dN/dS values in plastid genomes of *Rhopalocnemis* and *Balanophora* are lower with statistical significance than in photosynthetic relatives (Su et al., 2019; Schelkunov, Nuraliev & Logacheva, 2019). Therefore, the negative selection is likely stronger in *Rhopalocnemis* and *Balanophora*, not weaker. Just as the increase in the AT content, the increase in the substitution rate without relaxation of negative selection is supposed to be a typical feature of highly reduced plastid genomes of non-photosynthetic plants, not only the ones from the family Balanophoraceae (Logacheva, Schelkunov & Penin, 2011; Schelkunov et al., 2015; Lam, Soto Gomez & Graham, 2015; Logacheva et al., 2016; Roquet et al., 2016; Braukmann et al., 2017). In less reduced plastid genomes that still experience loss of photosynthesis-related genes, selection acting on the genes being lost is relaxed (Barrett & Davis, 2012; Wicke et al., 2013). We must warn that while the calculation of the AT content and its comparison between species is straightforward, the calculation of substitution rates and selection is less reliable. The main reason is the saturation on long branches which may lead to underestimation of substitution rates and, also, to underestimation of dN/dS (dos Reis & Yang, 2013). The underestimation of dN/dS on long branches follows from the fact that non-synonymous positions reach saturation faster than synonymous. Therefore, the seemingly not increased dN/dS values in plastid genomes of nonphotosynthetic species may be a computational artefact that is a consequence of the increased substitution rate.

The analysis for nuclear genes shows that in nuclear genes of *Rhopalocnemis phalloides* and *Balanophora fungosa*, just as in plastid ones, substitutions occur faster than in genes of photosynthetic relatives (Fig. 3a). At the same time, dN/dS values for nuclear genes of *Rhopalocnemis phalloides* and *Balanophora fungosa* are lower than in photosynthetic relatives (Fig 3b, p-value < 10^−5^ by the likelihood ratio test). This means a stronger negative selection. Or, alternatively, despite the low p-value this may be a consequence of the computational problem described above.

**Figure 3.**
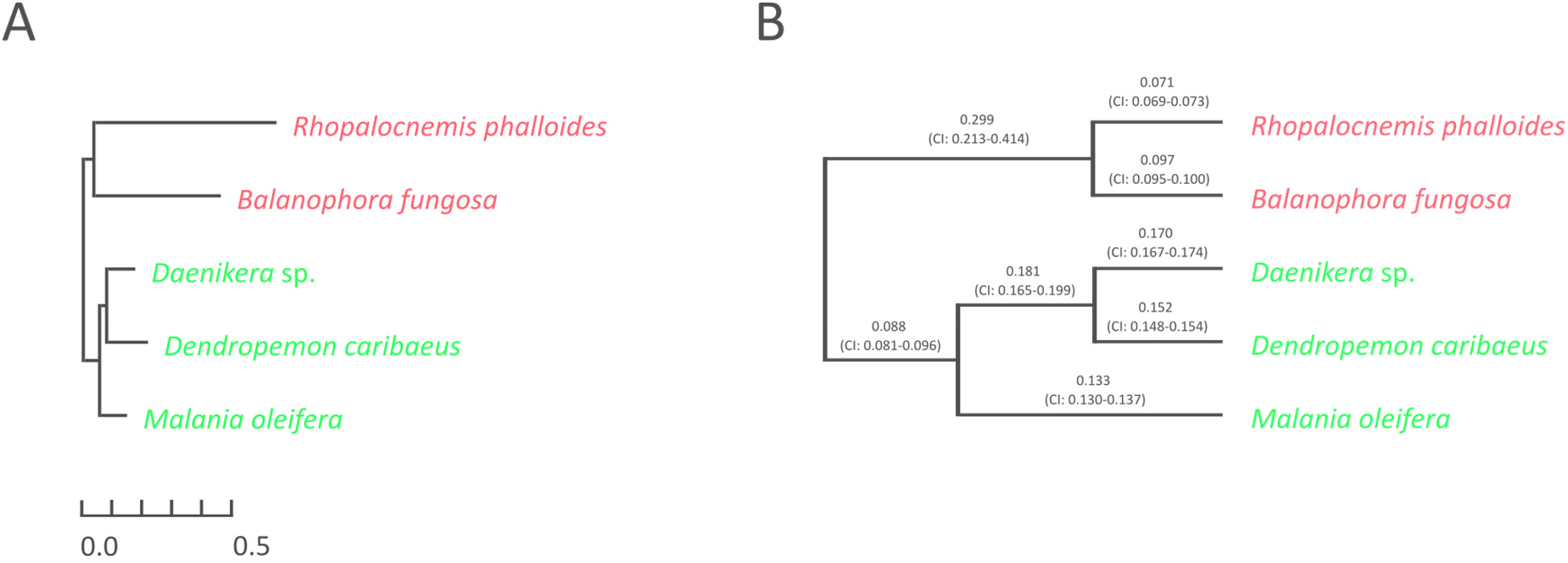
Evolutionary parameters of nuclear genes in the studied Santalales. (A) The phylogram where branch lengths represent numbers of substitutions per position. (B) The cladogram that depicts dN/dS values on branches with their 95% confidence intervals (CIs). *Arabidopsis thaliana*, used as the outgroup, is not shown. Names of non-photosynthetic species are in red, while names of photosynthetic species are in green. All bootstrap support values for the topology are 100%.

The dN/dS ratio on the branch of the common ancestor of *Rhopalocnemis phalloides* and *Balanophora fungosa* is, at the same time, significantly higher (p-value < 10^−5^ by the likelihood ratio test) than in photosynthetic plants. We are unaware of the cause of this increase in the dN/dS. Though there was likely a massive loss of photosynthesis-related genes on that branch accompanied by relaxation of selection in those genes, the analysis of dN/dS was performed using only extant genes, thus that relaxation of selection in the lost genes cannot be an explanation for this effect.

### The causes of unusual AT contents and substitution rates in the plastid genomes of Rhopalocnemis phalloides and Balanophora fungosa

As was described above, the plastid genomes of *Rhopalocnemis phalloides* and *Balanophora fungosa* have increased AT contents and substitution rates. Taking into account that all genes that encode proteins participating in plastid genome replication, recombination and repair (RRR) are encoded in the nuclear genome, the clue for these features of the plastid genome is to be sought in the nuclear one.

Based on a literature analysis we composed a list of 6 RRR proteins that were previously experimentally shown to function in RRR of the plastid genome, but not of mitochondrial or nuclear genomes (Table S3, part “Genes encoding other replication, recombination and repair proteins that work solely in plastids”). The RBH analysis of photosynthetic species indicates transcripts of 5 of these proteins present in *Daenikera* sp., 5 in *Dendropemon caribaeus* and 6 in *Malania oleifera*. However, we found transcripts of only 1 protein in *Rhopalocnemis phalloides* and only 2 in *Balanophora fungosa*. The transcripts absent in both *Rhopalocnemis phalloides* and *Balanophora fungosa* while present in photosynthetic species are of the following proteins:

1. MUTS2. The exact function of this protein in plants is unknown, but its bacterial homologs were shown to promote recombination suppression (Pinto et al., 2005).
2. OSB2. This protein also likely suppresses recombination (Zaegel et al., 2006).
3. RECA. It functions in the recombination-dependent DNA repair (Rowan, Oldenburg & Bendich, 2010). Additionally, *Rhopalocnemis phalloides* lacks the transcript of the following protein:
4. ARP. A nuclease which repairs DNA lesions coming from oxidative damage (Akishev et al., 2016).

The disappearance of plastid RRR proteins gives an explanation for why the plastid mutation accumulation rate is so high in *Rhopalocnemis phalloides* and *Balanophora fungosa*. Since gene conversion, a recombination-dependent process, is supposed to be GC-biased in plastids (Wu & Chaw, 2015; Zhitao Niu et al., 2017), disruption of recombination-dependent repair may make the genome more AT-rich, which may explain the high AT contents of the plastid genomes of *Rhopalocnemis phalloides* and *Balanophora fungosa*.

A transcriptomic analysis of non-photosynthetic plants of the genera *Epipogium* and *Hypopitys*, which also have AT-rich plastids with an increased rate of nucleotide substitutions, also demonstrated the absence of RECA, though MUTS2, OSB2 and ARP were present there (Schelkunov, Penin & Logacheva, 2018). This may point to the universality of the link between the high plastid AT content, high plastid substitution rates and loss of genes coding for RRR proteins.

An important question is why these genes are being lost in non-photosynthetic plants. The Accelerated Junk Removal (AJR) hypothesis proposed by us earlier (Schelkunov, Nuraliev & Logacheva, 2019) indicates that the increased mutation rate in the plastid genome may be beneficial for non-photosynthetic plants because it accelerates the removal of pseudogenes in the plastid genome. Indeed, after a plant loses the ability to photosynthesise, selection does not act on photosynthesis-related genes anymore. They start to accumulate deleterious mutations and their products may become dangerous to the plastids. Therefore, an increase in the mutation accumulation rate may be beneficial since it accelerates the complete disappearance of a gene or at least disappearance of genomic elements like promoters of ribosomal binding sites required for expression of the gene. Though scientists usually study the rate of nucleotide substitutions in plastids of non-photosynthetic plants, the rates of accumulation of indels and structural mutations were also shown to be increased there (Wicke et al., 2016; Wicke & Naumann, 2018).

Analyses of substitution rates in plastids of different lineages of non-photosynthetic plants indicated that the substitution rate in increased not only on the branch where photosynthesis was lost but also on branches of descendants (Schelkunov et al., 2015; Feng et al., 2016; Braukmann et al., 2017; Schelkunov, Nuraliev & Logacheva, 2019; Chen et al., 2020a). If the AJR hypothesis is correct, the explanation for this may be the irreversibility of the RRR gene loss. The mutation accumulation rate will be high until new RRR genes with plastid-targeted products evolve to replace the lost ones.

An alternative to AJR may be a hypothesis that postulates that the harm from a loss of an RRR gene is roughly proportional to the number of genes in which the mutation accumulation rate will increase after that RRR gene is lost. Simply speaking, imagine a 100,000 bp-long genome and a protein which fixes 100 errors after each replication of this genome. If the genome shortens to 10,000 bp, then after each replication this protein will fix only 10 errors. Therefore, the existence of this protein becomes less beneficial for the organism, even if strong negative selection acts on genes remaining in this small genome. This implies that when the plastid genome of a non-photosynthetic plant loses photosynthesis-related genes, the selection acting on the plastid genome RRR machinery relaxes, leading to the loss of RRR genes. This, in turn, may lead to the increase in the substitution rate and the AT content in the remaining plastid genes. We further call this the Less Important Accuracy (LIA) hypothesis. A similar explanation for the negative correlation between genome sizes and their mutation rates was proposed earlier by Drake et al. (1998).

Of the aforementioned 6 RRR proteins that function in plastids, a transcript of only 1 protein was found in both *Rhopalocnemis phalloides* and *Balanophora fungosa*, namely, of NTH1, which functions in the base excision repair (Roldán-Arjona et al., 2000). Its dN/dS on the branches of the studied non-photosynthetic plants is 0.20, while on the branches of the studied photosynthetic plants it is 0.22, with the p-value for the difference, as calculated by the likelihood ratio test, being 0.92. A similar analysis conducted previously for retained RRR proteins that function in plastids of *Epipogium* and *Hypopitys* also demonstrates that dN/dS values of their genes do not differ significantly from that of photosynthetic relatives (Schelkunov, Penin & Logacheva, 2018). Thus, selection on those RRR proteins that have survived does not seem to be relaxed.

The increased substitution rate of the nuclear genome (Fig. 3), in our opinion, may be explained in several ways:

1. The coevolution of a host and its parasite is often described as an “arms race” in which the host constantly invents mechanisms of defence, while the parasite in turn shall invent ways to avoid the defence. In such a situation, it may be beneficial both for the host and for the parasite to have increased mutation rates (Haraguchi & Sasaki, 1996). Since *Rhopalocnemis phalloides* and *Balanophora fungosa* rely on parasitism more than their mixotrophic relatives *Daenikera* sp., *Dendropemon caribaeus* and *Malania oleifera* that were used as photosynthetic species of comparison, it is logical that the arms race is more intense in *Rhopalocnemis phalloides* and *Balanophora fungosa*. Thus, the higher mutation rate and, consequently, the higher rate of substitutions (fixed point mutations) may theoretically be beneficial for *Rhopalocnemis phalloides* and *Balanophora fungosa*. This effect, however, can be used to explain the nuclear substitution rate but not the plastid substitution rate, because the plastid genome lacks genes whose products participate in the parasite-host interaction.
2. The AJR hypothesis can also explain the increased substitution rate in the nuclear genome, as photosynthetic plants contain a lot of genes with products functioning in photosynthesis (see Table S3, for example), thus the loss of photosynthesis and the subsequent degradation of these proteins is dangerous.
3. The LIA hypothesis may partially explain the increase in the nuclear substitution rate too. However, only a small fraction of nuclear genes of photosynthetic plants encodes proteins that participate in photosynthesis (Table 1). Consequently, the LIA effect is unlikely to be strong enough to explain the approximately threefold increase in the substitution rates in nuclear genes of *Rhopalocnemis phalloides* and *Balanophora fungosa*.

It was reported that the substitution rate is increased in the mitochondrial genomes of parasitic plants too (Bromham, Cowman & Lanfear, 2013). This may potentially be explained by the fact that there are RRR proteins common between the mitochondrial and the plastid genome, like RECA2 (Shedge et al., 2007), and thus the mitochondrial genome may also suffer if such proteins are lost or become less effective.

## Conclusions

The transcriptomic analysis of two non-photosynthetic plants from the family Balanophoraceae gives a hint to the cause of the extreme features of their plastid genomes. The likely cause is the loss of nuclear-encoded proteins that function in the plastid genome repair. However, *why* these proteins were lost remains enigmatic. The Accelerated Junk Removal hypothesis and the Less Important Accuracy hypothesis are two possible explanations.

## Acknowledgements

Sequencing on the HiSeq 4000 was performed using the resources of the Skoltech Genomics Core facility.

## References

Abascal F, Zardoya R, Telford MJ. 2010. TranslatorX: multiple alignment of nucleotide sequences guided by amino acid translations. Nucleic Acids Research 38:W7–W13. DOI: 10.1093/nar/gkq291.

Akishev Z, Taipakova S, Joldybayeva B, Zutterling C, Smekenov I, Ishchenko AA, Zharkov DO, Bissenbaev AK, Saparbaev M. 2016. The major Arabidopsis thaliana apurinic/apyrimidinic endonuclease, ARP is involved in the plant nucleotide incision repair pathway. DNA Repair 48:30–42. DOI: 10.1016/j.dnarep.2016.10.009.

Barrett CF, Davis JI. 2012. The plastid genome of the mycoheterotrophic *Corallorhiza striata* (Orchidaceae) is in the relatively early stages of degradation. American Journal of Botany 99:1513–1523. DOI: 10.3732/ajb.1200256.

Bellot S, Renner SS. 2015. The plastomes of two species in the endoparasite genus *Pilostyles* (Apodanthaceae) each retain just five or six possibly functional genes. Genome Biology and Evolution:evv251. DOI: 10.1093/gbe/evv251.

Berardini TZ, Reiser L, Li D, Mezheritsky Y, Muller R, Strait E, Huala E. 2015. The arabidopsis information resource: Making and mining the “gold standard” annotated reference plant genome: Tair: Making and Mining the “Gold Standard” Plant Genome. genesis 53:474–485. DOI: 10.1002/dvg.22877.

Bolger AM, Lohse M, Usadel B. 2014. Trimmomatic: a flexible trimmer for Illumina sequence data. Bioinformatics 30:2114–2120. DOI: 10.1093/bioinformatics/btu170.

Braukmann TWA, Broe MB, Stefanović S, Freudenstein JV. 2017. On the brink: the highly reduced plastomes of nonphotosynthetic Ericaceae. New Phytologist 216:254–266. DOI: 10.1111/nph.14681.

Bromham L, Cowman PF, Lanfear R. 2013. Parasitic plants have increased rates of molecular evolution across all three genomes. BMC evolutionary biology 13:126.

Buchfink B, Xie C, Huson DH. 2015. Fast and sensitive protein alignment using DIAMOND. Nature Methods 12:59–60. DOI: 10.1038/nmeth.3176.

Camacho C, Coulouris G, Avagyan V, Ma N, Papadopoulos J, Bealer K, Madden TL. 2009. BLAST+: architecture and applications. BMC Bioinformatics 10:421. DOI: 10.1186/1471-2105-10-421.

Caraballo-Ortiz MA, González-Castro A, Yang S, dePamphilis CW, Carlo TA. 2017. Dissecting the contributions of dispersal and host properties to the local abundance of a tropical mistletoe. Journal of Ecology 105:1657–1667. DOI: 10.1111/1365-2745.12795.

Carpenter EJ, Matasci N, Ayyampalayam S, Wu S, Sun J, Yu J, Jimenez Vieira FR, Bowler C, Dorrell RG, Gitzendanner MA, Li L, Du W, K. Ullrich K, Wickett NJ, Barkmann TJ, Barker MS, Leebens-Mack JH, Wong GK-S. 2019. Access to RNA-sequencing data from 1,173 plant species: The 1000 Plant transcriptomes initiative (1KP). GigaScience 8. DOI: 10.1093/gigascience/giz126.

Castresana J. 2000. Selection of conserved blocks from multiple alignments for their use in phylogenetic analysis. Molecular Biology and Evolution 17:540–552.

Chen X, Fang D, Wu C, Liu B, Liu Y, Sahu SK, Song B, Yang S, Yang T, Wei J, Wang X, Zhang W, Xu Q, Wang H, Yuan L, Liao X, Chen L, Chen Z, Yuan F, Chang Y, Lu L, Yang H, Wang J, Xu X, Liu X, Wicke S, Liu H. 2020a. Comparative Plastome Analysis of Root- and Stem-Feeding Parasites of Santalales Untangle the Footprints of Feeding Mode and Lifestyle Transitions. Genome Biology and Evolution 12:3663–3676. DOI: 10.1093/gbe/evz271.

Chen J, Yu R, Dai J, Liu Y, Zhou R. 2020b. The loss of photosynthesis pathway and genomic locations of the lost plastid genes in a holoparasitic plant Aeginetia indica. BMC Plant Biology 20. DOI: 10.1186/s12870-020-02415-2.

Cummings MP, Welschmeyer NA. 1998. Pigment composition of putatively achlorophyllous angiosperms. Plant Systematics and Evolution 210:105–111. DOI: 10.1007/BF00984730.

Drake JW, Charlesworth B, Charlesworth D, Crow JF. 1998. Rates of spontaneous mutation. Genetics 148:1667–1686.

El-Gebali S, Mistry J, Bateman A, Eddy SR, Luciani A, Potter SC, Qureshi M, Richardson LJ, Salazar GA, Smart A, Sonnhammer ELL, Hirsh L, Paladin L, Piovesan D, Tosatto SCE, Finn RD. 2019. The Pfam protein families database in 2019. Nucleic Acids Research 47:D427–D432. DOI: 10.1093/nar/gky995.

Emms DM, Kelly S. 2019. OrthoFinder: phylogenetic orthology inference for comparative genomics. Genome Biology 20. DOI: 10.1186/s13059-019-1832-y.

Feng Y-L, Wicke S, Li J-W, Han Y, Lin C-S, Li D-Z, Zhou T-T, Huang W-C, Huang L-Q, Jin X-H. 2016. Lineage-Specific Reductions of Plastid Genomes in an Orchid Tribe with Partially and Fully Mycoheterotrophic Species. Genome Biology and Evolution 8:2164–2175. DOI: 10.1093/gbe/evw144.

The Gene Ontology Consortium. 2019. The Gene Ontology Resource: 20 years and still GOing strong. Nucleic Acids Research 47:D330–D338. DOI: 10.1093/nar/gky1055.

Haas BJ, Papanicolaou A, Yassour M, Grabherr M, Blood PD, Bowden J, Couger MB, Eccles D, Li B, Lieber M, MacManes MD, Ott M, Orvis J, Pochet N, Strozzi F, Weeks N, Westerman R, William T, Dewey CN, Henschel R, LeDuc RD, Friedman N, Regev A. 2013. De novo transcript sequence reconstruction from RNA-seq using the Trinity platform for reference generation and analysis. Nature Protocols 8:1494–1512. DOI: 10.1038/nprot.2013.084.

Haraguchi Y, Sasaki A. 1996. Host–Parasite Arms Race in Mutation Modifications: Indefinite Escalation Despite a Heavy Load? Journal of Theoretical Biology 183:121–137. DOI: 10.1006/jtbi.1996.9999.

Kanehisa M, Sato Y, Kawashima M, Furumichi M, Tanabe M. 2016. KEGG as a reference resource for gene and protein annotation. Nucleic Acids Research 44:D457–462. DOI: 10.1093/nar/gkv1070.

Kanehisa M, Sato Y, Morishima K. 2016. BlastKOALA and GhostKOALA: KEGG Tools for Functional Characterization of Genome and Metagenome Sequences. Journal of Molecular Biology 428:726–731. DOI: 10.1016/j.jmb.2015.11.006.

Katoh K, Standley DM. 2013. MAFFT multiple sequence alignment software version 7: improvements in performance and usability. Molecular Biology and Evolution 30:772–780. DOI: 10.1093/molbev/mst010.

Klopfenstein DV, Zhang L, Pedersen BS, Ramírez F, Warwick Vesztrocy A, Naldi A, Mungall CJ, Yunes JM, Botvinnik O, Weigel M, Dampier W, Dessimoz C, Flick P, Tang H. 2018. GOATOOLS: A Python library for Gene Ontology analyses. Scientific Reports 8. DOI: 10.1038/s41598-018-28948-z.

Lam VKY, Soto Gomez M, Graham SW. 2015. The highly reduced plastome of mycoheterotrophic *Sciaphila* (Triuridaceae) is colinear with its green relatives and is under strong purifying selection. Genome Biology and Evolution 7:2220–2236. DOI: 10.1093/gbe/evv134.

Leake JR. 1994. The biology of myco-heterotrophic (‘saprophytic’) plants. New Phytologist 127:171–216. DOI: 10.1111/j.1469-8137.1994.tb04272.x.

Lee X-W, Mat-Isa M-N, Mohd-Elias N-A, Aizat-Juhari MA, Goh H-H, Dear PH, Chow K-S, Haji Adam J, Mohamed R, Firdaus-Raih M, Wan K-L. 2016. Perigone Lobe Transcriptome Analysis Provides Insights into Rafflesia cantleyi Flower Development. PLOS ONE 11:e0167958. DOI: 10.1371/journal.pone.0167958.

Li A-R, Mao P, Li Y-J. 2019. Root hemiparasitism in Malania oleifera (Olacaceae), a neglected aspect in research of the highly valued tree species. Plant Diversity 41:347–351. DOI: 10.1016/j.pld.2019.09.003.

Lim GS, Barrett CF, Pang C-C, Davis JI. 2016. Drastic reduction of plastome size in the mycoheterotrophic Thismia tentaculata relative to that of its autotrophic relative Tacca chantrieri. American Journal of Botany 103:1129–1137. DOI: 10.3732/ajb.1600042.

Liu Y, Popp B, Schmidt B. 2014. CUSHAW3: Sensitive and Accurate Base-Space and Color-Space Short-Read Alignment with Hybrid Seeding. PLoS ONE 9:e86869. DOI: 10.1371/journal.pone.0086869.

Logacheva MD, Schelkunov MI, Penin AA. 2011. Sequencing and analysis of plastid genome in mycoheterotrophic orchid *Neottia nidus-avis*. Genome Biology and Evolution 3:1296–1303. DOI: 10.1093/gbe/evr102.

Logacheva MD, Schelkunov MI, Shtratnikova VY, Matveeva MV, Penin AA. 2016. Comparative analysis of plastid genomes of non-photosynthetic Ericaceae and their photosynthetic relatives. Scientific Reports 6:30042. DOI: 10.1038/srep30042.

Marcin J, Julita M, José C, May M, Marc-André S, Etienne D. 2020. The genomic impact of mycoheterotrophy: targeted gene losses but extensive expression reprogramming. Plant Biology. DOI: 10.1101/2020.06.26.173617.

Merckx V, Bidartondo MI, Hynson NA. 2009. Myco-heterotrophy: when fungi host plants. Annals of Botany 104:1255–1261. DOI: 10.1093/aob/mcp235.

Mistry J, Finn RD, Eddy SR, Bateman A, Punta M. 2013. Challenges in homology search: HMMER3 and convergent evolution of coiled-coil regions. Nucleic Acids Research 41:e121–e121. DOI: 10.1093/nar/gkt263.

Molina J, Hazzouri KM, Nickrent D, Geisler M, Meyer RS, Pentony MM, Flowers JM, Pelser P, Barcelona J, Inovejas SA, Uy I, Yuan W, Wilkins O, Michel C-I, LockLear S, Concepcion GP, Purugganan MD. 2014. Possible loss of the chloroplast genome in the parasitic flowering plant *Rafflesia lagascae (Rafflesiaceae)*. Molecular Biology and Evolution 31:793–803. DOI: 10.1093/molbev/msu051.

Mower JP, Jain K, Hepburn NJ. 2012. The Role of Horizontal Transfer in Shaping the Plant Mitochondrial Genome. In: Advances in Botanical Research. Elsevier, 41–69. DOI: 10.1016/B978-0-12-394279-1.00003-X.

Naumann J, Der JP, Wafula EK, Jones SS, Wagner ST, Honaas LA, Ralph PE, Bolin JF, Maass E, Neinhuis C, Wanke S, dePamphilis CW. 2016. Detecting and characterizing the highly divergent plastid genome of the nonphotosynthetic parasitic plant *Hydnora visseri* (Hydnoraceae). Genome Biology and Evolution 8:345–363. DOI: 10.1093/gbe/evv256.

Ng S-M, Lee X-W, Mat-Isa M-N, Aizat-Juhari MA, Adam JH, Mohamed R, Wan K-L, Firdaus-Raih M. 2018. Comparative analysis of nucleus-encoded plastid-targeting proteins in Rafflesia cantleyi against photosynthetic and non-photosynthetic representatives reveals orthologous systems with potentially divergent functions. Scientific Reports 8. DOI: 10.1038/s41598-018-35173-1.

Nickrent DL. 2020. Parasitic angiosperms: How often and how many? TAXON 69:5–27. DOI: 10.1002/tax.12195.

Park J, Suh Y, Kim S. 2020. A complete chloroplast genome sequence of *Gastrodia elata* (Orchidaceae) represents high sequence variation in the species. Mitochondrial DNA Part B 5:517–519. DOI: 10.1080/23802359.2019.1710588.

Patro R, Duggal G, Love MI, Irizarry RA, Kingsford C. 2017. Salmon provides fast and bias-aware quantification of transcript expression. Nature Methods 14:417–419. DOI: 10.1038/nmeth.4197.

Pinto AV, Mathieu A, Marsin S, Veaute X, Ielpi L, Labigne A, Radicella JP. 2005. Suppression of Homologous and Homeologous Recombination by the Bacterial MutS2 Protein. Molecular Cell 17:113–120. DOI: 10.1016/j.molcel.2004.11.035.

dos Reis M, Yang Z. 2013. Why Do More Divergent Sequences Produce Smaller Nonsynonymous/Synonymous Rate Ratios in Pairwise Sequence Comparisons? Genetics 195:195–204. DOI: 10.1534/genetics.113.152025.

Roldán-Arjona T, García-Ortiz M-V, Ruiz-Rubio M, Ariza RR. 2000. cDNA cloning, expression and functional characterization of an Arabidopsis thaliana homologue of the Escherichia coli DNA repair enzyme endonuclease III. Plant Molecular Biology 44:43–52. DOI: 10.1023/A:1006429114451.

Roquet C, Coissac É, Cruaud C, Boleda M, Boyer F, Alberti A, Gielly L, Taberlet P, Thuiller W, Van Es J, Lavergne S. 2016. Understanding the evolution of holoparasitic plants: the complete plastid genome of the holoparasite *Cytinus hypocistis* (Cytinaceae). Annals of Botany 118:885–896. DOI: 10.1093/aob/mcw135.

Roulet ME, Garcia LE, Gandini CL, Sato H, Ponce G, Sanchez-Puerta MV. 2020. Multichromosomal structure and foreign tracts in the Ombrophytum subterraneum (Balanophoraceae) mitochondrial genome. Plant Molecular Biology 103:623–638. DOI: 10.1007/s11103-020-01014-x.

Rowan BA, Oldenburg DJ, Bendich AJ. 2010. RecA maintains the integrity of chloroplast DNA molecules in Arabidopsis. Journal of Experimental Botany 61:2575–2588. DOI: 10.1093/jxb/erq088.

Sanchez-Puerta MV, Edera A, Gandini CL, Williams AV, Howell KA, Nevill PG, Small I. 2019. Genome-scale transfer of mitochondrial DNA from legume hosts to the holoparasite Lophophytum mirabile (Balanophoraceae). Molecular Phylogenetics and Evolution 132:243–250. DOI: 10.1016/j.ympev.2018.12.006.

Sanchez-Puerta MV, García LE, Wohlfeiler J, Ceriotti LF. 2017. Unparalleled replacement of native mitochondrial genes by foreign homologs in a holoparasitic plant. New Phytologist 214:376–387. DOI: 10.1111/nph.14361.

Schelkunov MI, Nuraliev MS, Logacheva MD. 2019. *Rhopalocnemis phalloides* has one of the most reduced and mutated plastid genomes known. PeerJ 7:e7500. DOI: 10.7717/peerj.7500.

Schelkunov MI, Penin AA, Logacheva MD. 2018. RNA-seq highlights parallel and contrasting patterns in the evolution of the nuclear genome of fully mycoheterotrophic plants. BMC Genomics 19. DOI: 10.1186/s12864-018-4968-3.

Schelkunov MI, Shtratnikova VY, Nuraliev MS, Selosse M-A, Penin AA, Logacheva MD. 2015. Exploring the limits for reduction of plastid genomes: a case study of the mycoheterotrophic orchids *Epipogium aphyllum* and *Epipogium roseum*. Genome Biology and Evolution 7:1179–1191. DOI: 10.1093/gbe/evv019.

Seregin A. 2018. Moscow University Herbarium (MW). DOI: 10.15468/cpnhcc.

Shedge V, Arrieta-Montiel M, Christensen AC, Mackenzie SA. 2007. Plant Mitochondrial Recombination Surveillance Requires Unusual RecA and MutS Homologs. THE PLANT CELL ONLINE 19:1251–1264. DOI: 10.1105/tpc.106.048355.

Simão FA, Waterhouse RM, Ioannidis P, Kriventseva EV, Zdobnov EM. 2015. BUSCO: assessing genome assembly and annotation completeness with single-copy orthologs. Bioinformatics 31:3210–3212. DOI: 10.1093/bioinformatics/btv351.

Stamatakis A. 2014. RAxML version 8: a tool for phylogenetic analysis and post-analysis of large phylogenies. Bioinformatics 30:1312–1313. DOI: 10.1093/bioinformatics/btu033.

Stöver BC, Müller KF. 2010. TreeGraph 2: Combining and visualizing evidence from different phylogenetic analyses. BMC Bioinformatics 11:7. DOI: 10.1186/1471-2105-11-7.

Su H-J, Barkman TJ, Hao W, Jones SS, Naumann J, Skippington E, Wafula EK, Hu J-M, Palmer JD, dePamphilis CW. 2019. Novel genetic code and record-setting AT-richness in the highly reduced plastid genome of the holoparasitic plant Balanophora. Proceedings of the National Academy of Sciences of the United States of America 116:934–943. DOI: 10.1073/pnas.1816822116.

Těšitel J. 2016. Functional biology of parasitic plants: a review. Plant Ecology and Evolution 149:5–20. DOI: 10.5091/plecevo.2016.1097.

Těšitel J, Těšitelová T, Minasiewicz J, Selosse M-A. 2018. Mixotrophy in Land Plants: Why To Stay Green? Trends in Plant Science 23:656–659. DOI: 10.1016/j.tplants.2018.05.010.

The Angiosperm Phylogeny Group. 2016. An update of the Angiosperm Phylogeny Group classification for the orders and families of flowering plants: APG IV. Botanical Journal of the Linnean Society 181:1–20. DOI: 10.1111/boj.12385.

Törönen P, Medlar A, Holm L. 2018. PANNZER2: a rapid functional annotation web server. Nucleic Acids Research 46:W84–W88. DOI: 10.1093/nar/gky350.

Vidal-Russell R, Nickrent DL. 2008. The first mistletoes: Origins of aerial parasitism in Santalales. Molecular Phylogenetics and Evolution 47:523–537. DOI: 10.1016/j.ympev.2008.01.016.

Wicke S, Müller KF, dePamphilis CW, Quandt D, Bellot S, Schneeweiss GM. 2016. Mechanistic model of evolutionary rate variation en route to a nonphotosynthetic lifestyle in plants. Proceedings of the National Academy of Sciences 113:9045–9050. DOI: 10.1073/pnas.1607576113.

Wicke S, Muller KF, de Pamphilis CW, Quandt D, Wickett NJ, Zhang Y, Renner SS, Schneeweiss GM. 2013. Mechanisms of functional and physical genome reduction in photosynthetic and nonphotosynthetic parasitic plants of the Broomrape Family. The Plant Cell 25:3711–3725. DOI: 10.1105/tpc.113.113373.

Wicke S, Naumann J. 2018. Molecular Evolution of Plastid Genomes in Parasitic Flowering Plants. In: Advances in Botanical Research. Elsevier, 315–347. DOI: 10.1016/bs.abr.2017.11.014.

Wickett NJ, Honaas LA, Wafula EK, Das M, Huang K, Wu B, Landherr L, Timko MP, Yoder J, Westwood JH, dePamphilis CW. 2011. Transcriptomes of the Parasitic Plant Family Orobanchaceae Reveal Surprising Conservation of Chlorophyll Synthesis. Current Biology 21:2098–2104. DOI: 10.1016/j.cub.2011.11.011.

Wu C-S, Chaw S-M. 2015. Evolutionary Stasis in Cycad Plastomes and the First Case of Plastome GC-Biased Gene Conversion. Genome Biology and Evolution 7:2000–2009. DOI: 10.1093/gbe/evv125.

Xu C-Q, Liu H, Zhou S-S, Zhang D-X, Zhao W, Wang S, Chen F, Sun Y-Q, Nie S, Jia K-H, Jiao S-Q, Zhang R-G, Yun Q-Z, Guan W, Wang X, Gao Q, Bennetzen JL, Maghuly F, Porth I, Van de Peer Y, Wang X-R, Ma Y, Mao J-F. 2019. Genome sequence of *Malania oleifera*, a tree with great value for nervonic acid production. GigaScience 8. DOI: 10.1093/gigascience/giy164.

Yang Z. 2007. PAML 4: phylogenetic analysis by maximum likelihood. Molecular Biology and Evolution 24:1586–1591. DOI: 10.1093/molbev/msm088.

Yang Y, He S-L. 2019. The complete chloroplast genome of *Malania oleifera* (Olacaceae), an endangered species in China. Mitochondrial DNA Part B 4:1867–1868. DOI: 10.1080/23802359.2019.1614493.

Yuan Y, Jin X, Liu J, Zhao X, Zhou J, Wang X, Wang D, Lai C, Xu W, Huang J, Zha L, Liu D, Ma X, Wang L, Zhou M, Jiang Z, Meng H, Peng H, Liang Y, Li R, Jiang C, Zhao Y, Nan T, Jin Y, Zhan Z, Yang J, Jiang W, Huang L. 2018. The Gastrodia elata genome provides insights into plant adaptation to heterotrophy. Nature Communications 9. DOI: 10.1038/s41467-018-03423-5.

Zaegel V, Guermann B, Le Ret M, Andrés C, Meyer D, Erhardt M, Canaday J, Gualberto JM, Imbault P. 2006. The Plant-Specific ssDNA Binding Protein OSB1 Is Involved in the Stoichiometric Transmission of Mitochondrial DNA in Arabidopsis. The Plant Cell 18:3548–3563. DOI: 10.1105/tpc.106.042028.

Zhitao Niu, Qingyun Xue, Hui Wang, Xuezhu Xie, Shuying Zhu, Wei Liu, Xiaoyu Ding. 2017. Mutational Biases and GC-Biased Gene Conversion Affect GC Content in the Plastomes of Dendrobium Genus. International Journal of Molecular Sciences 18:2307. DOI: 10.3390/ijms18112307.

Zhou N, Jiang Y, Bergquist TR, Lee AJ, Kacsoh BZ, Crocker AW, Lewis KA, Georghiou G, Nguyen HN, Hamid MN, Davis L, Dogan T, Atalay V, Rifaioglu AS, Dalkiran A, Cetin Atalay R, Zhang C, Hurto RL, Freddolino PL, Zhang Y, Bhat P, Supek F, Fernández JM, Gemovic B, Perovic VR, Davidović RS, Sumonja N, Veljkovic N, Asgari E, Mofrad MRK, Profiti G, Savojardo C, Martelli PL, Casadio R, Boecker F, Schoof H, Kahanda I, Thurlby N, McHardy AC, Renaux A, Saidi R, Gough J, Freitas AA, Antczak M, Fabris F, Wass MN, Hou J, Cheng J, Wang Z, Romero AE, Paccanaro A, Yang H, Goldberg T, Zhao C, Holm L, Törönen P, Medlar AJ, Zosa E, Borukhov I, Novikov I, Wilkins A, Lichtarge O, Chi P-H, Tseng W-C, Linial M, Rose PW, Dessimoz C, Vidulin V, Dzeroski S, Sillitoe I, Das S, Lees JG, Jones DT, Wan C, Cozzetto D, Fa R, Torres M, Warwick Vesztrocy A, Rodriguez JM, Tress ML, Frasca M, Notaro M, Grossi G, Petrini A, Re M, Valentini G, Mesiti M, Roche DB, Reeb J, Ritchie DW, Aridhi S, Alborzi SZ, Devignes M-D, Koo DCE, Bonneau R, Gligorijević V, Barot M, Fang H, Toppo S, Lavezzo E, Falda M, Berselli M, Tosatto SCE, Carraro M, Piovesan D, Ur Rehman H, Mao Q, Zhang S, Vucetic S, Black GS, Jo D, Suh E, Dayton JB, Larsen DJ, Omdahl AR, McGuffin LJ, Brackenridge DA, Babbitt PC, Yunes JM, Fontana P, Zhang F, Zhu S, You R, Zhang Z, Dai S, Yao S, Tian W, Cao R, Chandler C, Amezola M, Johnson D, Chang J-M, Liao W-H, Liu Y-W, Pascarelli S, Frank Y, Hoehndorf R, Kulmanov M, Boudellioua I, Politano G, Di Carlo S, Benso A, Hakala K, Ginter F, Mehryary F, Kaewphan S, Björne J, Moen H, Tolvanen MEE, Salakoski T, Kihara D, Jain A, Šmuc T, Altenhoff A, Ben-Hur A, Rost B, Brenner SE, Orengo CA, Jeffery CJ, Bosco G, Hogan DA, Martin MJ, O’Donovan C, Mooney SD, Greene CS, Radivojac P, Friedberg I. 2019. The CAFA challenge reports improved protein function prediction and new functional annotations for hundreds of genes through experimental screens. Genome Biology 20. DOI: 10.1186/s13059-019-1835-8.

