## Supplementary Figures for "Genomic comparison of non-photosynthetic plants from the family Balanophoraceae with their photosynthetic relatives"

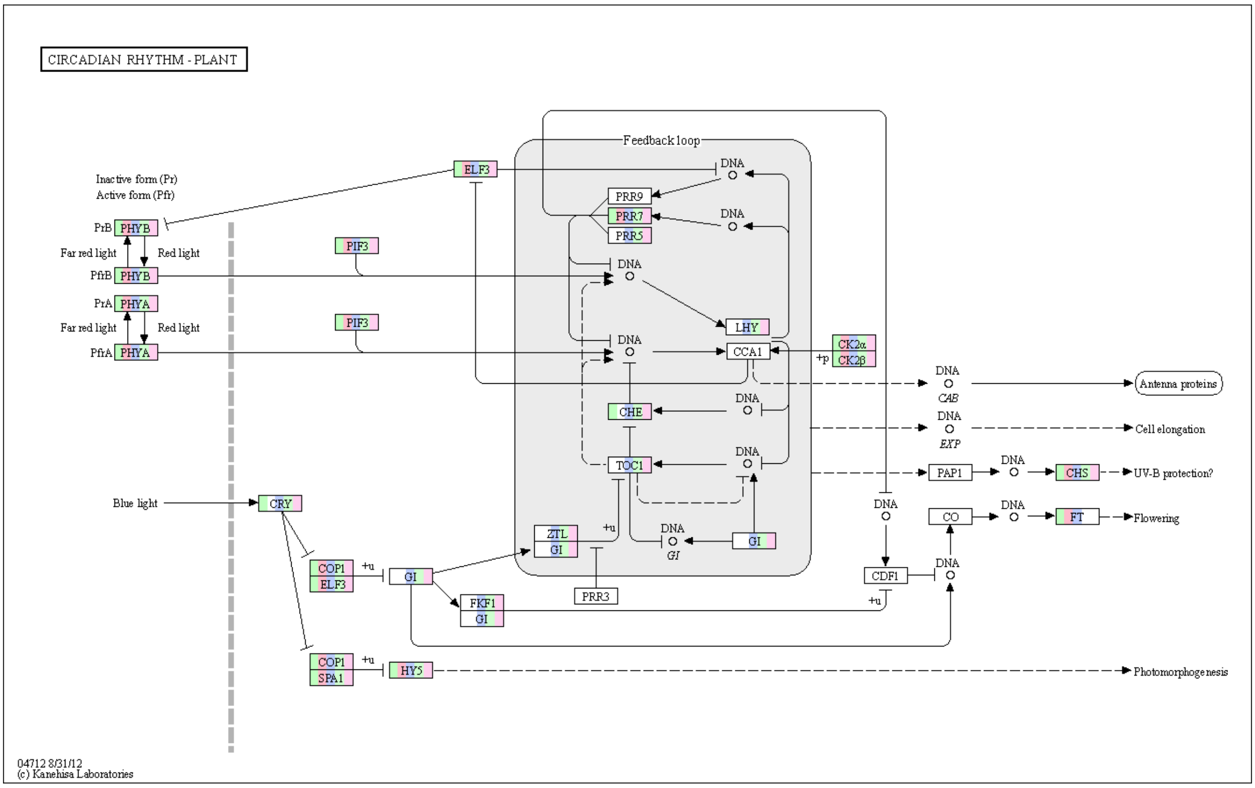


Figure S1. Proteins functioning in the circadian rhythm organisation in Santalales.

This map is a coloured metabolic map "Circadian rhythm - plant" from the KEGG database. Rectangles are proteins, circles are metabolites, lines are reactions, dashed lines are poorly studied reactions. The presence of colours in the rectangles, from left to right, denotes the presence of a transcript of the corresponding protein in: *Rhopalocnemis phalloides*, *Balanophora fungosa*, *Daenikera* sp., *Dendropemon caribaeus*, *Malania oleifera*. For example, the transcript of the protein GI is likely absent in *Rhopalocnemis phalloides* and *Balanophora fungosa*, but present in *Daenikera* sp., *Dendropemon caribaeus*, *Malania oleifera*.

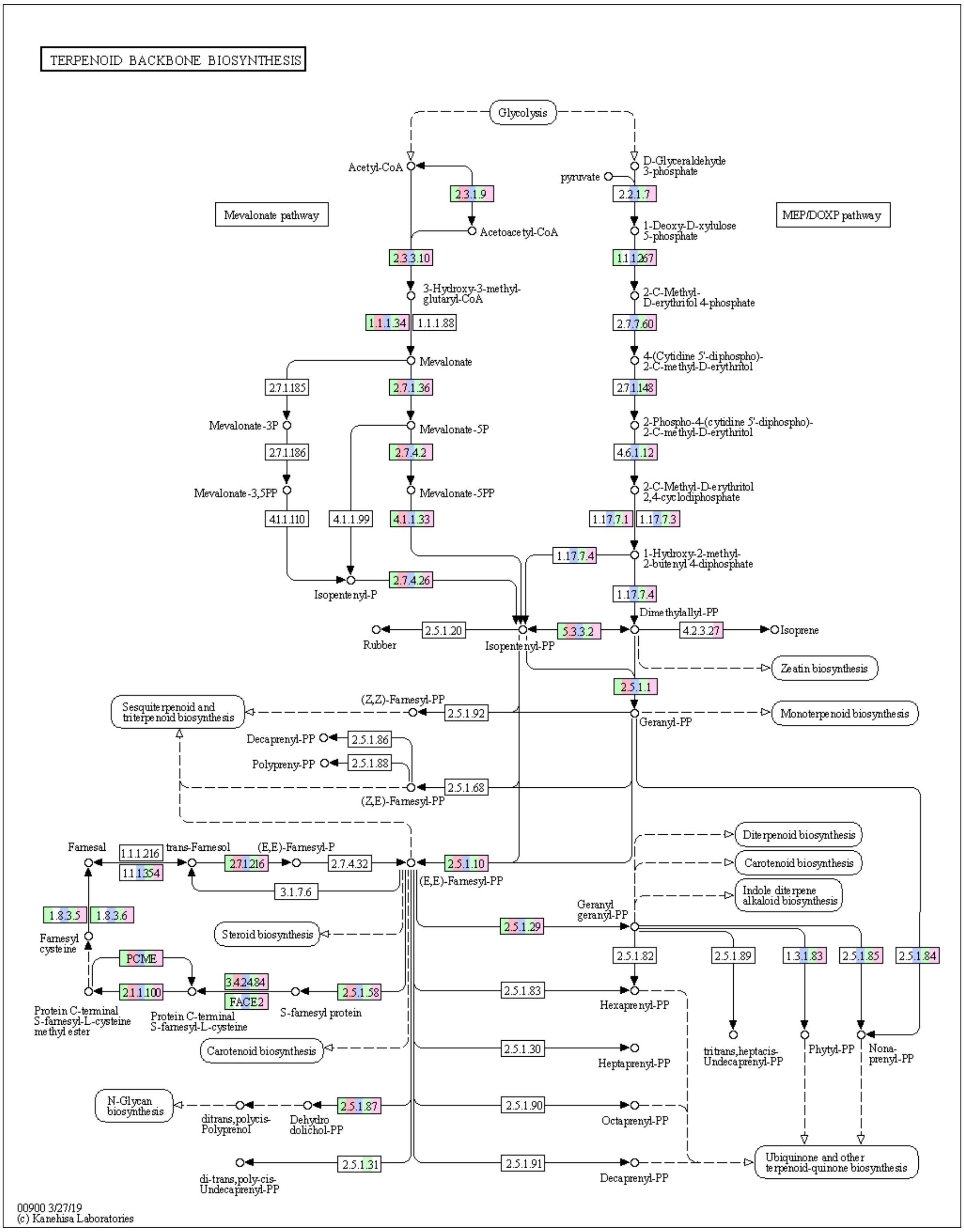


Figure S2. Proteins functioning in the terpenoid backbone synthesis in Santalales.

This map is a coloured metabolic map "Terpenoid backbone synthesis" from the KEGG database. Rectangles are proteins, circles are metabolites, lines are reactions, dashed lines are poorly studied reactions. Numbers in the rectangles are Enzyme Commission (EC) numbers, words in the rectangles are protein symbols. The presence of colours in the rectangles, from left to right, denotes the presence of a transcript of the corresponding protein in: *Rhopalocnemis phalloides*, *Balanophora fungosa*, *Daenikera* sp., *Dendropemon caribaeus*, *Malania oleifera*. For example, the transcript of the protein with the EC code 2.7.1.148 is likely absent in *Rhopalocnemis phalloides* and *Balanophora fungosa*, but present in *Daenikera* sp., *Dendropemon caribaeus*, *Malania oleifera*.

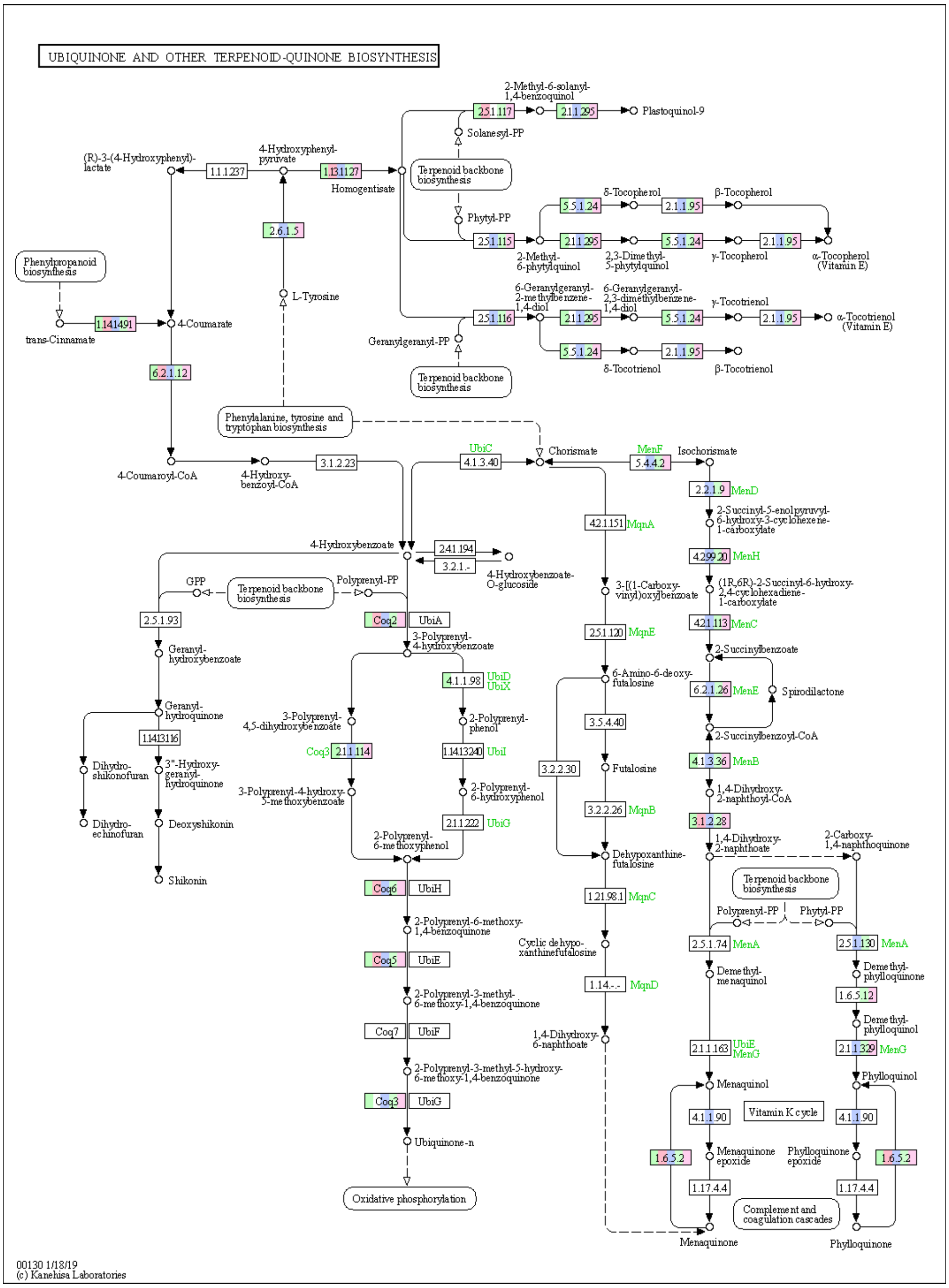


Figure S3. Proteins functioning in the ubiquinone and other terpenoid-quinone biosynthesis in Santalales.

This map is a coloured metabolic map "Ubiquinone and other terpenoid-quinone biosynthesis" from the KEGG database. Rectangles are proteins, circles are metabolites, lines are reactions, dashed lines are poorly studied reactions. Numbers in the rectangles are Enzyme Commission (EC) numbers, words in the rectangles or near them are protein symbols. The presence of colours in the rectangles, from left to right, denotes the presence of a transcript of the corresponding protein in: *Rhopalocnemis phalloides*, *Balanophora fungosa*, *Daenikera* sp., *Dendropemon caribaeus*, *Malania oleifera*. For example, the transcript of the protein with the EC code 4.2.1.113 is likely absent in *Rhopalocnemis phalloides* and *Balanophora fungosa*, but present in *Daenikera* sp., *Dendropemon caribaeus*, *Malania oleifera*.

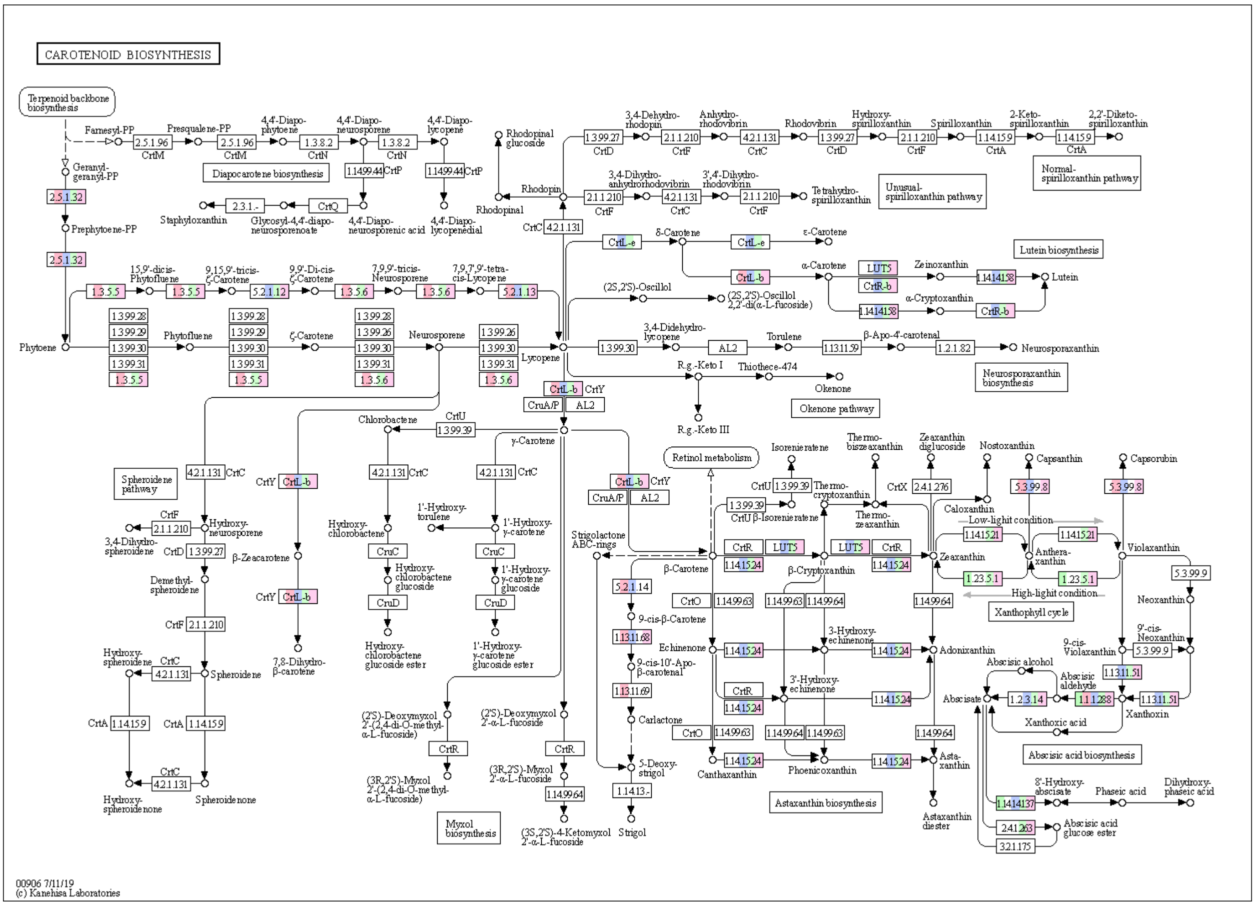


Figure S4. Proteins functioning in the carotenoid biosynthesis in Santalales.

This map is a coloured metabolic map "Carotenoid biosynthesis" from the KEGG database. Rectangles are proteins, circles are metabolites, lines are reactions, dashed lines are poorly studied reactions. Numbers in the rectangles are Enzyme Commission (EC) numbers, words in the rectangles or near them are protein symbols. The presence of colours in the rectangles, from left to right, denotes the presence of a transcript of the corresponding protein in: *Rhopalocnemis phalloides*, *Balanophora fungosa*, *Daenikera* sp., *Dendropemon caribaeus*, *Malania oleifera*. For example, the transcript of the protein LUT5 is likely absent in *Rhopalocnemis phalloides* and *Balanophora fungosa*, but present in *Daenikera* sp., *Dendropemon caribaeus*, *Malania oleifera*.
